# Autism associated *Cntnap2* deletion disrupts vestibular sensory signaling and spatial cognition in mice

**DOI:** 10.64898/2026.05.28.728446

**Authors:** Yvette Shu, Yushan Chen, Dyllan Zhou, Xiaoya Deng, Liliana D. Florea, Tara Deemyad, Soroush G. Sadeghi

## Abstract

Autism spectrum disorder (ASD) is frequently accompanied by sensory and motor abnormalities, including impaired balance, postural control, and spatial orientation, that are often attributed largely to altered central circuitry. Emerging evidence, however, suggests that peripheral sensory dysfunction can also shape ASD related behavioral phenotypes. Here, we tested whether loss of the ASD associated gene *Cntnap2*/Caspr2 alters vestibular signaling in *Cntnap2^-/-^* mice. Developmental transcriptomic analysis showed that *Cntnap2* is expressed in vestibular sensory organs and increases during the first postnatal month, coincident with vestibular pathway maturation. Vestibular sensory evoked potentials revealed reduced response amplitudes and prolonged latencies in *Cntnap2*^-/-^ mice, indicating impaired peripheral afferent responses to transient linear acceleration. *Cntnap2^-/-^* mice also showed delayed contact righting, reduced ocular counter roll, and increased hindlimb slips and compensatory tail excursions during balance beam walking, whereas rotational vestibulo-ocular reflex gain and phase were preserved. These vestibular and balance abnormalities were accompanied by reduced novel arm preference in the Y maze and severe impairment of Barnes maze acquisition, consistent with impaired spatial learning. Together, these findings identify *Cntnap2*/Caspr2 as a regulator of vestibular sensory signaling and support a model in which disrupted peripheral vestibular input, likely acting together with central effects of *Cntnap2* loss, contributes to sensorimotor and spatial cognitive phenotypes relevant to ASD.

## Introduction

Vestibular and balance problems are highly prevalent in individuals with autism spectrum disorder (ASD), with studies indicating that up to 80% of children with ASD exhibit vestibular or balance dysfunction [1, 2]. Studies of pediatric patients with ASD have demonstrated abnormalities in Sensory Organization Tests, videonystagmography, and rotary chair testing [1]. These clinically important features contribute to motor coordination challenges, postural instability, atypical sensory processing, and delays in developmental milestones such as sitting and walking [3]. Vestibular abnormalities in ASD are generally attributed to altered sensory integration within cortical and cerebellar circuits [2, 4]. Similarly, impairments in spatial navigation and memory have been documented in individuals with ASD and in animal models of ASD and are often attributed to abnormalities in the structure and function of the hippocampus [5–7]. However, recent studies have challenged a predominantly central circuit focused view of ASD by showing that dysfunction restricted to peripheral sensory neurons can directly produce behavioral phenotypes associated with the disorder. Selective disruption of peripheral somatosensory neurons is sufficient to produce tactile hypersensitivity and anxiety like behaviors in different mouse models of ASD [8–10], and the prevalence of ASD is higher in individuals with visual and auditory impairments [11–14]. These findings raise the possibility that altered peripheral sensory input may contribute to ASD related behaviors by shaping the development and function of central neural circuits.

The vestibular system is therefore a particularly relevant peripheral sensory pathway for investigating ASD related sensorimotor and spatial cognitive phenotypes because it provides the brain with continuous information about head motion, gravitational orientation, postural state, and self-motion. Vestibular organs in the inner ear detect angular and linear head acceleration and transmit this information through vestibular nerve afferents to brainstem and cerebellar circuits that regulate posture, balance, and reflexive eye movements. These signals are integrated with visual and proprioceptive inputs to support gaze stabilization, postural control, spatial orientation, and navigation [15–17]. Vestibular afferents comprise functionally distinct populations that encode complementary aspects of head motion and orientation. Regular afferents provide sustained signals related to head orientation relative to gravity [18] and contribute to the vestibulo ocular reflex (VOR) [19], whereas irregular afferents encode rapid head movements (Goldberg, 2000; Sadeghi et al., 2007; Ono et al., and may contribute to vestibulospinal reflexes [20, 21]. Vestibular signals encoding head motion and orientation also provide critical information to hippocampal and parahippocampal circuits involved in spatial learning and memory [15, 22–25]. Consistent with this role, patients with vestibular disorders frequently exhibit deficits in spatial memory and navigation [15, 23, 26, 27] and experimental disruption of vestibular input impairs hippocampal place cell activity and degrades spatial learning in rodents [15, 24, 28–30]. Therefore, vestibular dysfunction could provide a mechanistic link between balance abnormalities and spatial-cognitive deficits reported in ASD. However, the contribution of peripheral vestibular input from the inner ear to balance and spatial cognitive deficits in ASD remains poorly understood.

Genetic studies have identified contactin-associated protein-like 2 (*CNTNAP2*) as a prominent ASD associated neurodevelopmental risk gene [31, 32]. The mouse ortholog, *Cntnap2*, encodes Cntnap2 or Caspr2 [33], and *Cntnap2*^-/-^ mice are widely used as a genetic model relevant to ASD. These animals show ASD related phenotypes, including social deficits, repetitive behaviors, altered neural connectivity, and seizures [34, 35]. Previous studies have also reported impairments in motor coordination and spatial memory in individuals or mice carrying CNTNAP2/*Cntnap2* mutations [5, 34, 35]. In addition, anti-CNTNAP2 antibodies can produce autoimmune encephalopathy that is commonly accompanied by dizziness [36, 37], further suggesting that CNTNAP2/Caspr2 related mechanisms may affect vestibular or balance related function.

Several lines of evidence suggest that loss of *Cntnap2* may disrupt vestibular signaling at the level of peripheral sensory organs. First, Caspr2 is expressed in the vestibular periphery [38, 39]. Caspr2 is a membrane associated protein that regulates synaptic organization, ion channel localization, neural excitability, and myelination [33, 40, 41], all of which are important for sensory signal transmission. Second, studies of mice lacking Caspr1, a related contactin associated protein, show structural abnormalities at the specialized synapse between type I vestibular hair cells and calyx afferent terminals, including enlargement of the synaptic cleft and altered localization of KCNQ potassium channels in the calyx terminal [42]. These changes may disrupt potassium dependent nonquantal transmission between type I hair cells and calyx terminals [43–49], a specialized form of signaling that enables rapid encoding of head movement [45, 48, 50] and contributes to vestibular reflexes such as the VOR [48, 51]. Finally, because Caspr2 influences synaptic organization and myelination [40, 52–54], signaling at glutamatergic synapses between type II vestibular hair cells and bouton afferents, as well as conduction along vestibular afferents, could also be affected. Nevertheless, whether ASD associated genes such as *Cntnap2* regulate vestibular sensory function, and whether disruption of vestibular input contributes to balance and spatial cognitive deficits in the absence of *Cntnap2*, has not been systematically investigated.

Here, we used *Cntnap2*^-/-^ mice to test the hypothesis that Caspr2 is required for normal peripheral vestibular signaling and that its loss is associated with vestibular dependent sensorimotor deficits and impaired spatial learning and memory. We combined developmental transcriptomic analysis of vestibular sensory organs with vestibular sensory evoked potentials, gravity dependent reflex assays, ocular motor testing, quantitative balance beam analysis, and hippocampus dependent spatial memory tasks. We show that *Cntnap2* is expressed in vestibular sensory organs during postnatal development and that loss of Caspr2 reduces vestibular nerve responses, impairs gravity dependent reflexes and balance behavior, and is associated with deficits in spatial working memory and spatial learning. These findings identify *Cntnap2*/Caspr2 as a regulator of vestibular sensory signaling and support a model in which disrupted vestibular input may contribute to sensorimotor and spatial cognitive phenotypes relevant to ASD.

## Materials and Methods

### Animals

All procedures were approved by the Johns Hopkins University Institutional Animal Care and Use Committee and followed National Institutes of Health guidelines. WT C57BL/6J (strain 000664) and Cntnap2 mutant (strain 017482) mice were obtained from The Jackson Laboratory and bred in-house; offspring were used for all experiments, with WT littermates as controls when possible. Mice were housed under standard conditions (12 h light/dark and ad libitum food/water). Experiments were conducted during the light phase in 2–10 month old male and female mice, with investigators blinded to genotype and sex.

### RNA Sequencing

RNA sequencing and analysis were performed to assess gene expression in vestibular tissue across developmental stages. Whole tissue cristae and maculae were dissected from the bony labyrinth of the inner ear in mice at postnatal days 7, 14, 21, and 30. Total RNA was extracted using standard RNA isolation procedures. RNA quality was assessed prior to sequencing. Sequence reads were aligned to the mouse reference genome GRCm39 using STAR v2.7.11b [55] and gene expression levels were quantified using DESeq2 [56] against the GENCODE v.M28 mouse gene annotations. Differential expression analysis was performed using DESeq2, with adjusted p-values calculated to correct for multiple comparisons. Gene expression patterns were visualized using volcano plots, developmental expression profiles, and heatmaps of selected genes associated with neuronal connectivity and synaptic organization.

### Vestibular Sensory Evoked Potential (VsEP) recording

Short-latency VsEP recordings were used to assess vestibular nerve function as previously described (Jones et al., 2002, 2011; Raghu et al., 2020). Mice were anesthetized (ketamine 80 mg/kg, xylazine 10 mg/kg, i.p.) and subcutaneous electrodes were placed at the vertex (recording electrode), right ear, and left ear. With the head secured to an electromagnetic shaker (ET-132, Labworks Inc.), linear naso-occipital stimuli (2 ms; 0.5–2 g/ms, 3 dB steps) were delivered. Signals were amplified (×100,000) with a Grass P511 amplifier, filtered (0.3–3 kHz), and digitized at 100 kHz (micro1401, CED). Responses (250 trials/direction) were averaged, and peak-to-peak amplitude of the first positive and negative waveform (P1-N1) and P1 latency were measured using custom MATLAB scripts.

### Contact Righting Reflex

Contact righting reflex was used to assess vestibular-mediated detection of gravity, as previously described [48]. Mice were lightly anesthetized with isoflurane and positioned supine between two flat surfaces such that all four paws contacted the upper surface. Timing began upon recovery of consciousness, and latency to righting was recorded as the time required to assume a prone position, defined as at least three paws contacting the lower surface. Mice with dysfunction of peripheral vestibular sensors would remain supine and even walk upside-down for at least 30 s, whereas normal mice would flip over in a few seconds [48]. Surfaces were cleaned with 70% ethanol between trials.

### Rotational vestibulo-ocular reflex (VOR) and ocular counter roll (OCR)

VOR was recorded as described previously [48]. Headposts were implanted under isoflurane, and after 2 – 5 days recovery, mice were acclimated (3 sessions). Animals were restrained with the head fixed and the left eye was tracked by an infrared camera (ISCAN) during rotations in the dark (0.3 – 6 Hz; 20 – 45 °/s). Pilocarpine (1–2%) was applied to constrict pupils. Signals were acquired with a CED 1401 digitizer and analyzed in MATLAB. Eye velocity was derived from position, saccades removed, and gain/phase computed by the best fit of head velocity sines to eye velocity traces [57]. OCR was measured immediately after VOR. Mice were tilted ±10 ° and ±20 ° in the roll plane and held for 15 s in random order. OCR amplitude was the change in eye position from baseline to the end of the tilt time; gain was calculated as eye position change divided by tilt angle [58].

### Balance Beam

Dynamic balance and coordination were assessed using a narrow beam (10 mm × 60 cm) leading to a dark escape box. Mice were habituated (10 min), then completed three training crossings from a brightly lit start platform. After a 10 minute rest, three test trials were recorded (30 fps; posterior view) with 20 s intertrial rest. The apparatus was cleaned with 70% ethanol between trials. Analysis spanned from full paw contact on the beam to nose entry into the goal box. Crossing times were measured and hindlimb slips were counted by two independent experimenters and averaged. Trials were excluded if recordings were compromised by external factors or technical errors; consequently, 1–3 valid trials were available per mouse and used for analysis. Next, tail kinematics were quantified using DeepLabCut [59, 60]. A model (tail base, midpoint, tip) was trained on 400 annotated frames and iteratively refined to ensure high reliability; low-confidence detections were first excluded based on likelihood threshold of 0.6, and tracking outlier frames were removed using a 3×IQR filter applied to the radial distances from the tail base to the tail midpoint and tail tip. Custom Python scripts reconstructed tail position to derive tail angle and movement dynamics. Tail angle was then calculated from the filtered tail tip position relative to the tail base. Tail swing amplitude was quantified using a shortest arc approach to account for the circular nature of angular data and avoid artificial inflation when angles crossed the −180°/180° boundary.

### Y-Maze

Short-term spatial memory was assessed using a two-trial novelty preference Y-maze [61]. The maze had three identical arms (35 × 5 cm, 10 cm walls) arranged at 120° with arm specific visual cues. In the habituation phase, one arm was blocked and mice explored the other two arms for 15 min, followed by a 1-hour delay. In the test phase, all arms were open (the previously blocked arm served as novel) and mice explored for 5 min. Time spent per arm was recorded; preference for the novel arm indicated intact memory. The blocked arm was counterbalanced, and the maze was cleaned with 70% ethanol between trials.

### Barnes Maze

Spatial learning and memory were assessed using the Barnes maze [62, 63]. The apparatus was a 60 cm circular platform with 20 evenly spaced perimeter holes; one led to a dark escape box. Visual wall cues provided spatial guidance and bright lighting provided motivation for finding the escape hole. For each animal, there were habituation, acquisition, and probe phases. On Day 1, mice were placed in the escape box for habituation (1 min), then allowed to explore for up to 5 min or until entering the escape hole (guided if necessary). After 1 hour, acquisition began with a new, fixed escape box location. Mice completed two trials/day for 10 days (1 h intertrial interval; results averaged). Each trial started under a randomized start box location; mice had 3 min to find the escape hole or were guided and held in the escape box for ∼15 s. The maze was cleaned with 70% ethanol between trials. Latency to escape (time to enter and remain at least 5 s in the escape box) and path length traveled to escape hole were measured. A probe test was performed 3 days after the final acquisition session to assess memory retention. During the probe test, the escape box was removed, and mice were allowed to explore the maze for 1 min. Memory retention was assessed by tracking the time required to reach the previous escape location and the distance traveled before reaching that location. Sessions were recorded at 30 fps with an overhead infrared camera (White Matter LLC) and videos were analyzed using DeepLabCut [59, 60]. Pose estimation was performed using the DeepLabCut SuperAnimal top view mouse model as a pretrained model for transfer learning and fine tuning [64]. The model was fine tuned using 200 annotated frames and iteratively refined to ensure reliable tracking performance. Detections with likelihood < 0.6 were excluded. Distance travelled and latency to find the escape hole were quantified for each session with custom Python scripts.

### Statistical Analysis

Statistical analyses were conducted using GraphPad Prism (GraphPad Software, San Diego, CA, USA), R (R Foundation for Statistical Computing, Vienna, Austria), and Python. For uncensored data, normality was assessed using the Shapiro-Wilk test for sample sizes < 50 and the Kolmogorov-Smirnov test for sample sizes ≥ 50. Homogeneity of variance was evaluated using Levene’s test. Normally distributed data with equal variances were analyzed using parametric ANOVA models appropriate to the number of experimental factors. Where homogeneity of variance was not met, Welch’s ANOVA with Games-Howell post hoc testing was used. For data that did not satisfy normality assumptions, an aligned rank transform (ART) ANOVA was used, with post hoc pairwise comparisons performed using Wilcoxon tests with Holm correction. Comparisons against a fixed value, such as chance level, were performed using a one-sample t-test for normally distributed data or a Wilcoxon signed-rank test when normality assumptions were not met. For censored data from Barnes maze, comparisons were quantified by Cox proportional hazards regression to account for trials in which animals did not reach the defined endpoint within the allotted testing period. Data are presented as mean ± standard error of the mean (SEM) unless otherwise indicated. All tests were two sided and statistical significance was set at α = 0.05.

## Results

### Expression of *Cntnap2* in the vestibular neuroepithelium in the inner ear

Previous immunohistology studies have shown the expression of CNTNAP2/Caspr2 at the top of dimorphic calyx terminals (i.e., afferent terminals with both calyx and boutons) in adult rats [38, 39]. The vestibular periphery in the inner ear continues development during the 2-3 weeks after birth, as vestibular type I hair cells and their calyx terminal take form anatomically and develop their membrane properties [65, 66]. To determine the expression pattern of *Cntnap2* during the early development of the vestibular sensory organs in mouse inner ears, we analyzed RNA sequencing datasets collected weekly during the first postnatal month from a mix of cristae (from the semicircular canals) and maculae (of the otolith organs). Gene expression analysis revealed that *Cntnap2* transcripts were present in the vestibular tissue. Further, differential expression (DE) analysis showed statistically significant changes in the expression levels of several genes associated with neuronal connectivity and synaptic organization, including *Cntnap2*, *Shank3*, *Cacna1c*, *Syngap1* and *Nlgn3* (Fig. 1A). An adjusted p-value (padj) cutoff of 0.05 was considered for DE. Analysis of normalized read counts across developmental stages showed that *Cntnap2* expression varied across tested postnatal time points of P7, P14, P21, and P30. Notably, a heatmap clearly showed different patterns of variation for these genes, with the clustering of upregulated genes *Cntnap2*, *Kcnq3*, *Cacna2d3*, and *Shank2*, and the opposite pattern of downregulation for *Nlgn3, Cacna1c, Shank3, and Syngap1* genes (Fig. 1B). Furthermore, these results indicate that during the second half of the first postnatal month compared with the first two weeks after birth, these genes are part of a process that simultaneously upregulates *Cntnap2* (log_2_(fold change or FC) = 1.5) and *Cacna2d3* (log_2_FC = 2.0) and drives up the expression of *Kcnq3* (log_2_FC = 1.1) and *Shank2* (log_2_FC = 0.9) in a more subtle way. In contrast, the same period is associated with significant downregulation of *Shank3* (log_2_FC = - 1.6), *Cacna1c* (log_2_FC = -1.6), *Syngap1* (log_2_FC = -0.8) and *Nlgn3* (log_2_FC = -1.5). To better visualize temporal changes, we calculated fold changes in counts relative to P7 for these eight ASD associated genes (Fig. 1C). The resulting plots are in line with the heatmap and confirm the co-expression of *Cntnap2* and three other genes (*Kcnq3, Cacna2d3,* and *Shank2*) and, separately, of *Shank3*, *Nlgn3*, *Syngap1*, and *Cacna1c* (Fig. 1D). Together, the above findings provide strong evidence for the expression of *Cntnap2* in the inner ear, with an upregulation later in development.

**Figure 1.**
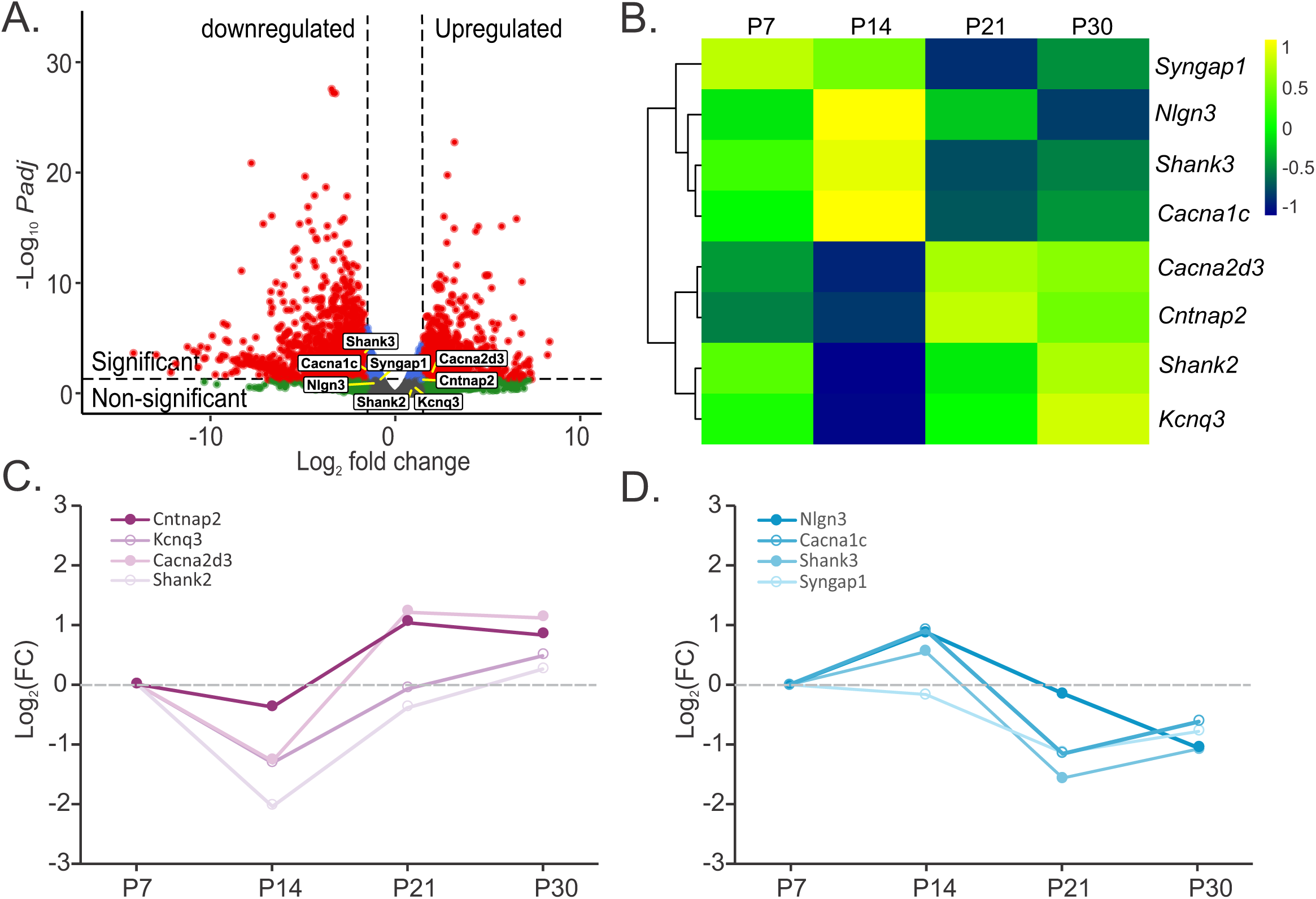
Expression of *Cntnap2* in the vestibular inner ear during the first postnatal month. **(A)** Volcano plot showing different genes expressed in the cristae and maculae of the vestibular organs in the inner ear. Several autism spectrum disorder (ASD) associated genes are highlighted. Horizontal dashed line shows p-adj = 0.05; vertical dashed lines are at -1.5 and 1.5 fold changes. **(B)** Heatmap of normalized read counts for eight selected ASD related genes across postnatal developmental stages (P7, P14, P21, and P30). **(C, D)** Temporal expression profiles (normalized changes relative to P7) of representative upregulated and downregulated genes, respectively, showing dynamic regulation during development.

### Impaired VsEP responses in *Cntnap2*^-/-^ mice

To determine whether Caspr2 encoded by *Cntnap2*, contributed to peripheral vestibular signaling, we recorded VsEP as a measure of the response of the vestibular nerve to fast linear head movements in *Cntnap2* mutant mice and wild type (WT) littermates at 2-3 months of age (Fig. 2A). Responses were quantified by measuring the peak-to-peak amplitude of the first positive (P1) and negative (N1) wave (P1–N1 amplitude) and P1 latency across stimulus intensities. Shapiro-Wilk tests indicated that response amplitudes were approximately normally distributed across groups. Representative traces show the reduced amplitude in a *Cntnap2^-/-^* mouse compared to a WT littermate (Fig. 2B). For all tested animals, *Cntnap2^-/-^* mice (n = 41) exhibited reduced waveform amplitudes compared to WT controls (n = 36) (two-way ANOVA, genotype main effect, F(1, 75) = 37.54, p < 0.0001) at all stimulus intensities tested (Post hoc Bonferroni test, p < 0.001 for all) (Fig. 2C). Comparison of P1 latencies between WT and *Cntnap2^-/-^* mice showed a significant difference at stimulus intensities of 0.75 and higher (mixed-effects aligned rank transform (ART) ANOVA, F(1, 75) = 39.99, p < 0.001 for genotype effect, Post hoc Wilcoxon test with Holm correction, p < 0.01 for all except 0.5 g/ms stimulus) (Fig. 2D). These results suggest an abnormal peripheral vestibular function in Cntnap2^-/-^ mice.

**Figure 2.**
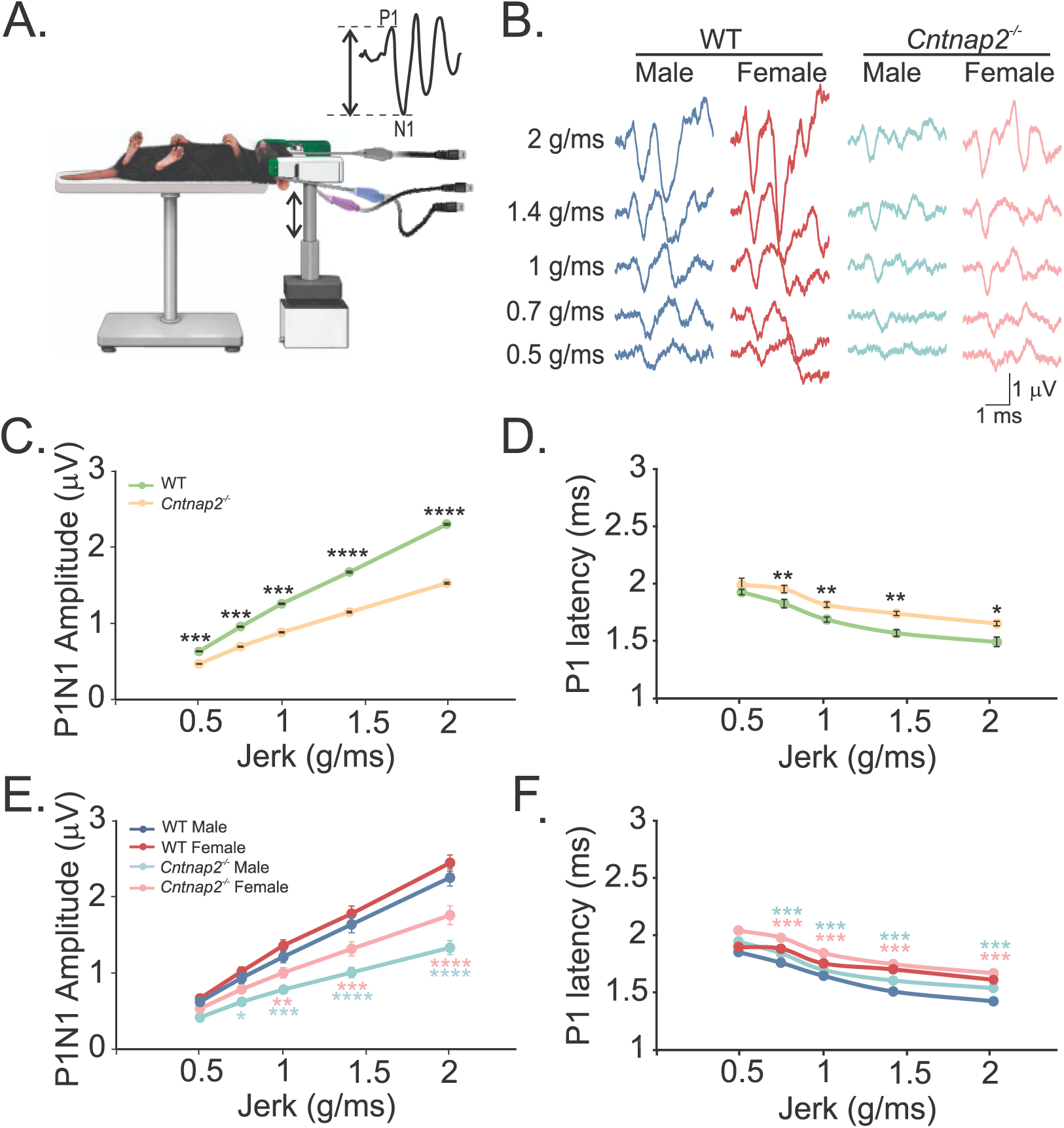
Dysfunction of the vestibular nerve in *Cntnap2*^-/-^ mice. **(A)** Schematic of VsEP recording apparatus and mouse positioning. Mice were anesthetized and placed in a prone position with subcutaneous electrodes positioned at the vertex and behind the ears (pinna). The head was secured to a mechanical shaker and subjected to repeated linear jerk stimuli in the naso-occipital (up-down) direction, activating both utricular and saccular vestibular pathways. Vestibular nerve responses were averaged, and peak-to-peak P1–N1 amplitudes and P1 latencies were quantified. **(B)** Representative VsEP traces from WT male, WT female, *Cntnap2^-/-^* male, and *Cntnap2^-/-^*female mice across increasing jerk intensities. WT mice exhibited larger P1–N1 amplitudes compared to *Cntnap2^-/-^* mice. **(C)** Average P1-N1amplitudes of VsEP responses show smaller responses in *Cntnap2^-/-^* mice compared to WT mice at all stimulus amplitudes. **(D)** Average P1 latencies of VsEP responses were longer in *Cntnap2^-/-^* mice compared to WT mice, except for the smallest stimulus amplitude. **(E)** Average amplitudes of WT male, WT female, *Cntnap2^-/-^*male, and *Cntnap2^-/-^* female vestibular nerve responses. A three-way repeated-measures ANOVA showed significant effects of stimulus amplitude (F(4, 292) = 594.87, p < 0.001), sex (F(1, 73) = 7.60, p = 0.007), and genotype (F(1, 73) = 42.59, p < 0.001), as well as a significant stimulus amplitude × genotype interaction (F(4, 292) = 31.54, p < 0.001). Post hoc Tukey tests showed reduced amplitudes in *Cntnap2^-/-^* males vs WT males at all intensities (p < 0.01 for all). In females, reductions were observed at 0.75 g/ms and higher (p < 0.01). **(F)** Average P1 latencies of WT male, WT female, *Cntnap2^-/-^* male, and *Cntnap2^-/-^* female vestibular nerve responses. Comparison by an ART ANOVA revealed significant main effects of stimulus intensity (F(4, 292) = 177.74, p < 0.0001), genotype (F(1, 73) = 36.93, p < 0.0001), and sex (F(1,73) = 4.12, p = 0.046), but no significant interactions (all p > 0.13). Post hoc Wilcoxon tests with Holm correction showed a strong overall genotype effect (p = 2×10⁻⁷), with significant differences between WT and *Cntnap2^-/-^* mice in both males and females (p < 0.001 for both). When examined across stimulus intensities, genotype differences were not significant at 0.5 g/ms (p = 0.089) but were significant from 0.75 g/ms to 2 g/ms (p < 0.001 for all). No significant sex differences were detected within either genotype (WT: p = 0.98; *Cntnap2^-/-^*: p = 0.84). All average data are presented as mean ± SEM. Statistical significances compared to WT are shown by * for p < 0.05, ** for p < 0.01, *** for p < 0.001, and **** for p < 0.0001.

Since autism phenotypes are generally stronger in males than females in most animal models and in individuals with ASD [67, 68], we also investigated male and female VsEP responses separately. For the population of recordings from the four group of mice (male and female, WT and *Cntnap2^-/-^*), P1–N1 amplitudes increased as a function of stimulus intensity across all groups (Fig. 2E). A three-way repeated-measures ANOVA revealed significant main effects of stimulus intensity (F(4, 292) = 594.87, p < 0.001), sex (F(1, 73) = 7.60, p = 0.007), and genotype (F(1, 73) = 42.59, p < 0.001), as well as a significant stimulus intensity × genotype interaction (F(4,292) = 31.54, p < 0.001), while other interactions were not significant. Tukey’s multiple comparisons tests showed significant differences between WT and KO responses at all stimulus intensities above 0.5 g/ms (p < 0.01 for all). The largest differences were at the highest intensity (2 g/ms) for all comparisons (p < 0.0001), indicating robust attenuation of responses in *Cntnap2^-/-^*mice at strong stimuli. Furthermore, while P1 latency decreased with increasing stimulus intensity across all groups, P1 latencies were significantly longer in *Cntnap2^-/-^* animals (Fig. 2F), with significant main effects of stimulus intensity, genotype, and sex (ART ANOVA, stimulus intensity: F(4, 292) = 177.74, p < 0.0001; genotype: F(1, 73) = 36.93, p < 0.0001; sex: F(1, 73) = 4.12, p = 0.046). There were no significant interaction effects. Post-hoc Wilcoxon tests with Holm correction revealed a strong overall genotype effect across both sexes (p < 0.00001) and no effect of sex across genotypes (WT: p = 0.98, *Cntnap2^-/-^*: p = 0.84). Note, the magnitude of latency differences remained small (in μs range), suggesting limited functional relevance.

Together, these findings show that loss of Caspr2 affects vestibular nerve responses to fast movements and provide clear evidence for a peripheral vestibular dysfunction in Cntnap2^-/-^ mice.

### Impaired gravity signaling in *Cntnap2*^-/-^ mice

It is believed that gravity is encoded by regular vestibular afferents [18] through quantal glutamate transmission [48]. To investigate whether altered peripheral vestibular function was associated with impaired gravity signaling, we used two behavioral tests. First, we assessed the contact righting reflex as a measure of gravity perception [48, 69]. When WT mice are put in a supine position between two surfaces, they flip over to a prone position in a few seconds [48] (Fig. 3B).

**Figure 3.**
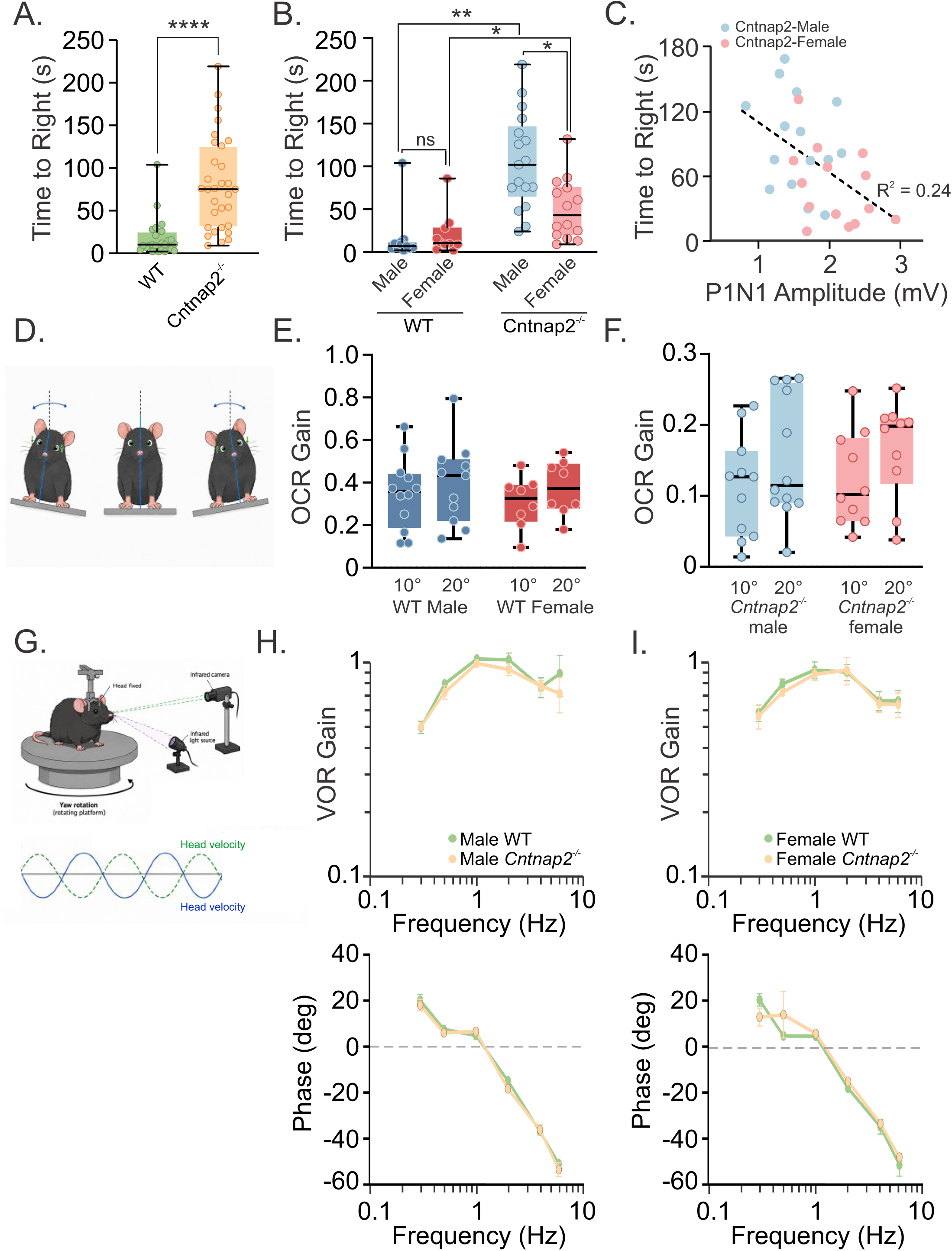
Detection of gravity is impaired in *Cntnap2*^-/-^ mice. **(A)** The time to turn from a supine position to the prone position (i.e., ‘time to right’) was increased in *Cntnap2^-/-^*mice compared to WT littermate. **(B)** Male and female *Cntnap2^-/-^* mice showed an increase in latency compared to their corresponding WT littermates. Note the wider range of latencies for male KO mice compared to female KO mice. **(C)** A fairly strong negative correlation is observed between responses of the contact righting reflex and VsEP amplitudes in *Cntnap2^-/-^* mice. Note the distribution of male and female data, with males having lower VsEP amplitudes and longer latencies. **(D)** Schematic of the OCR test. Mice were tilted in roll and compensatory change in vertical eye position was measured as an ocular readout of otolith dependent sense of gravity. **(E)** Box plots showing OCR gain in male and female WT mice during 10° and 20° roll-tilt stimuli. Black line shows the median and whiskers represent the range. **(F)** Box plots showing OCR gain in male and female *Cntnap2^-/-^*mice during 10° and 20° roll-tilt stimuli. Note, y-axis range compared to WT mice (panel E) and the significantly smaller OCR gains in *Cntnap2^-/-^* mice. These results support the observed poor sensation of gravity with righting reflex. **(G)** Schematic of the VOR setup. Head rotation was delivered on a turntable, in the dark while eye position was recorded by an infrared camera that moved with the animal. VOR gain and phase were calculated across stimulus frequencies. **(H)** VOR gain and phase were similar for male WT and *Cntnap2^-/-^* mice over the tested frequencies. **(I)** VOR gain and phase were similar for female WT and *Cntnap2^-/-^* mice over the tested frequencies. Data are plotted as mean ± SEM for VOR frequency response curves. Statistical comparisons are indicated in the figure: ns, not significant; * for p < 0.05; ** for p < 0.01; and *** for p < 0.0001.

In contrast, Cntnap2^-/-^ mice (n = 31) exhibited significantly longer righting latencies compared to WT mice (n = 20) (Mann-Whitney test, U = 81.50, p < 0.0001) (Fig. 3A). We then compared male and female mice latencies (ART ANOVA) and found a significant effect for both sex (F(1,47) = 13.14, p = 0.0007) and genotype (F(1,47) = 43.63, p < 0.0001). Post hoc pairwise Wilcoxon tests with Holm correction were then used to probe these effects. Within genotype comparisons showed a significant sex difference in Cntnap2^-/-^ mice (Post hoc Wilcoxon test with Holm correction, p = 0.01), but not in WT mice (p = 0.446). Within-sex comparisons showed significantly increased righting latencies in Cntnap2^-/-^ males compared to WT males (Post hoc Wilcoxon test with Holm correction, p < 0.0001), and in Cntnap2^-/-^ females compared to WT females (p = 0.005) (Fig. 3B).

To explore the relationship between vestibular nerve function and behavioral performance in Cntnap2^-/-^ mice, we correlated VsEP amplitudes with righting latencies from the same animals (Fig. 3C). These measures were negatively correlated (n = 29, r² = 0.2437; p = 0.0065 for slope), indicating that reduced vestibular responses were associated with slower righting reflexes. Together, these findings demonstrate impaired gravity perception associated with reduced peripheral vestibular function in Cntnap2^-/-^ mice.

Second, we measured the gain of the ocular counter roll (OCR), which is a reflex that uses changes in the gravity signal during head tilts to stabilize the gaze by trying to bring the eyes back to the horizontal line (Fig. 3D). We found that both male (n = 6) and female (n = 5) *Cntnap2*^-/-^ mice showed lower gains with decreases of up to 80% compared to male (n = 6) and female (n = 4) WT littermates in response to 10 ° and 20 ° tilts to right and left (Figs. 3E and 3F) (ANOVA, Tilt x Genotype interaction, males: F(3, 28) = 12.41, p < 0.0001; females: F(3, 21) = 9.92, p = 0.0003). There were no within genotype differences (males: F(1, 10) = 0.81, p = 0.39; females: F(1, 7) = 0.0022, p = 0.96).

These results suggest impaired detection of gravity in the absence of Caspr2, which could be due to an impaired glutamate transmission in the vestibular periphery [48]. However, we could not rule out the presence of coexistent deficits in central pathways due to lack of Caspr2 expression.

### Normal VOR responses in *Cntnap2*^-/-^ mice

We next tested VOR responses, which are believed to depend on NQ transmission between type I hair cells and calyx terminals [48, 51] of regular afferents [19]. Eye movements were recorded in alert head restrained mice rotated in darkness across different frequencies (Fig. 3G). To quantify the response, the gain and phase of eye velocities were calculated relative to velocities of table rotation at each frequency (0.3 – 6 Hz and 20 – 45 °/s). At any tested frequency, there was no difference between Cntnap2^-/-^ mice (6 male and 5 female) and WT littermates (6 males and 4 females) in VOR gains (ANOVA, Frequency x Genotype interaction, males: F(5, 49) = 0.49, p = 0.78; females: F(5, 41) = 0.068, p = 1.00) and phases (males: F(5, 49) = 1.10, p = 0.37; females: F(5, 40) = 0.94, p = 0.47) (Figs. 3H and 3I). These normal VOR results in Cntnap2^-/-^ mice suggest that lack of Caspr2 does not affect the NQ signaling between type I vestibular hair cells and calyx afferent terminals for regular afferents. This also suggests a lack of impairment in the central vestibulo-ocular pathway, suggesting that the observed OCR deficiencies are also most likely due to peripheral causes. However, we recognize that we cannot rule out peripheral dysfunction along with central compensation in the VOR pathway.

### Abnormal balance in *Cntnap2*^-/-^ mice

We next assessed the function of the vestibulospinal pathways using the balance beam test. Here, mice crossed a narrow (10 cm wide) elevated beam from an open start platform to an enclosed goal platform, a task requiring coordination and postural control that depends on vestibulospinal pathways and gravity sensation (i.e., sense of being vertical). For this test, we used 7 WT males, 8 WT females, 8 KO males and 9 KO females. Each animal was tested 3 times and results were averaged. Interestingly, crossing times were similar between Cntnap2^-/-^ and WT mice (two-way ANOVA, F (1, 28) = 0.7376, p = 0.40) and between sexes (F (1, 28) = 1.957, p = 0.17) and no significant interaction between sex and genotype was observed (F (1, 28) = 0.1546, p = 0.70). No significant differences were observed between male and female mice: WT male mice crossed in 9.2 ± 0.6 s and KO male mice crossed in 9.57 ± 0.8 s; WT female mice crossed in 7.3 ± 0.5 s and KO female mice crossed in 8.2 ± 0.6 s (Fig. 4A).

**Figure 4.**
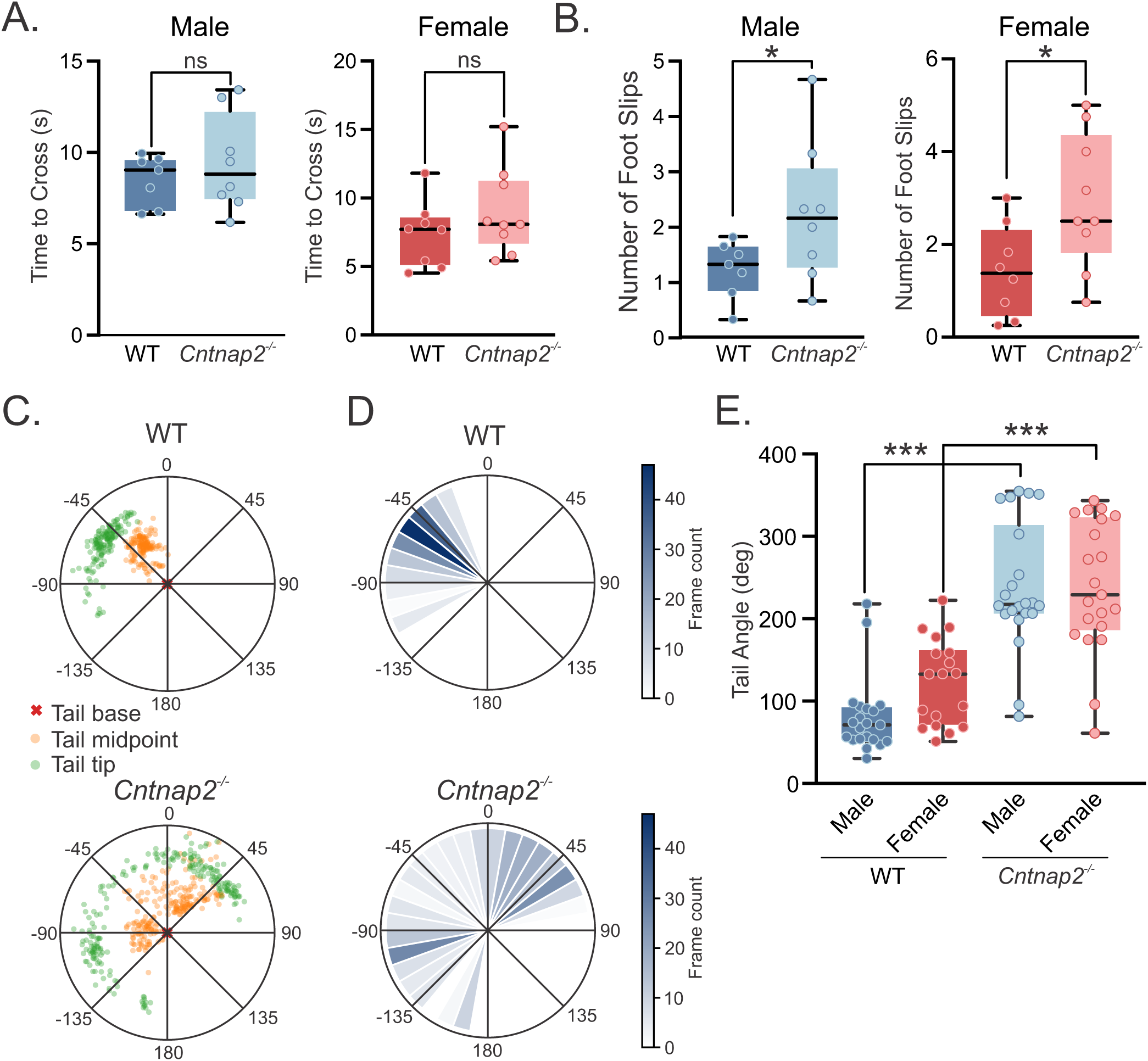
Abnormal dynamic balance is uncovered in *Cntnap2*^-/-^ mice using balance beam test. **(A)** No significant differences in crossing time were detected between WT and *Cntnap2^-/-^*mice in either males or females. **(B)** When the number of hind-limb foot slips were considered during beam crossing, *Cntnap2^-/-^* mice made significantly more foot slips than WT littermates (ART ANOVA, genotype effect, *p* = 0.00025, post-hoc Wilcoxon tests with Holm’s correction, *p* = 0.029 for male WT vs *Cntnap2^-/-^* and *p* = 0.012 for female WT and *Cntnap2^-/-^*). There was no significant effect of sex (p = 0.48) or sex × genotype interaction (p = 0.34). **(C)** Representative tail-tracking outputs from WT and *Cntnap2^-/-^*male mice during balance beam crossing. Tail position was reconstructed from markerless pose estimation of the tail base (red cross), tail midpoint (orange), and tail tip (green), plotted relative to the beam axis. Note broader angular tail movements in the *Cntnap2^-/-^* example compared to the WT mouse. **(D)** Representative plot showing frame-by-frame tail-angle distributions for WT and *Cntnap2^-/-^* mice. The increased spread in the *Cntnap2^-/-^* mouse illustrates larger lateral tail excursions during beam crossing, consistent with compensatory tail movements used to maintain balance. **(E)** Summary of tail-angle measurements in WT and *Cntnap2^-/-^* males and females. Male and female *Cntnap2^-/-^* mice exhibited significantly larger tail angles than WT littermates (ART ANOVA; genotype effect, p < 0.00001; Post-hoc Wilcoxon tests with Holm’s correction, p < 0.00001 for WT vs. *Cntnap2^-/-^* males and for WT and *Cntnap2^-/-^* females). There was no effect of sex (p = 0.10) or sex × genotype interaction (p = 0.13).

To further quantify the videos, we used two more measures. First, two independent observers counted the number of hind limb slips in the video recordings and results were averaged. Male WT and Cntnap2^-/-^ mice experienced 1.2 ± 0.2 and 2.3 ± 0.4 slips, respectively and female WT and Cntnap2^-/-^ mice experienced 1.3 ± 0.3 and 2.9 ± 0.4 slips, respectively (two-way ANOVA, significant main effect of genotype (F(1, 28) = 9.46, p = 0.0046), but no significant effect of sex (F(1, 28) = 1.11, p = 0.302) or sex × genotype interaction (F(1, 28) = 0.35, p = 0.562) (Fig. 4B).

Second, we compared tail positions between the different groups. Recent studies have identified the tail as an important contributor during locomotion in mice, particularly under conditions that challenge postural stability [70, 71]. These studies showed that mice swing their tails to counteract roll perturbations, generating angular momentum to stabilize themselves. We quantified tail movements using a DeepLabCut model to track the frame-by-frame position of the base, midpoint, and tip of the tail. Figure 4C shows a polar plot of representative examples of frame-by-frame positions of tail midpoint and tip for a WT and a Cntnap2^-/-^ mouse data, corrected for tail base position (i.e., overlapping tail bases for each frame). Notice that while crossing the beam, the tip of the tail of the WT mouse stayed mainly in the 0° – 90° range on the left side, but for the KO mouse it moved between the right and left sides and reached up to 180°, meaning that it touched the beam. Furthermore, the tip of the tail of the WT mouse was most of the time in the 45° to the left position, while the KO mouse tail was most of the time located about 90° to the left or 45° to the right (Fig. 4D). For quantitative population analysis, maximum tail angle provided the best metric for distinction between WT and KO mice. Two-way ANOVA revealed a significant main effect of genotype on maximum tail angle (F(1,78) = 88.86, p < 0.0001), with no significant main effect of sex (F(1,78) = 2.06, p = 0.155) and no genotype × sex interaction (F(1,78) = 2.05, p = 0.156). Cntnap2^−/−^ mice showed markedly larger maximum tail angles compared to WT controls (WT: 100.36 ± 8.50°, n = 39; Cntnap2^−/−^: 238.73 ± 11.80°, n = 43), indicating broader tail excursions during beam crossing (Fig. 4E).

Together, these findings indicate that loss of Caspr2 impairs balance beam performance and may reflect deficits in vestibulospinal pathways.

### Impaired spatial working memory in *Cntnap2*^-/-^ mice

Previous studies have shown that vestibular inputs to the hippocampus provide critical information about head motion and orientation necessary for the formation of spatial representations [15, 22–26, 29, 30]. Disruption of vestibular signaling should impair hippocampus dependent spatial learning and memory. Given the vestibular deficits observed in Cntnap2^-/-^ mice, we next investigated whether these animals also exhibit impairments in spatial memory tasks. We assessed spatial working (short-term) memory using the two-trial novelty preference Y-maze task, in which animals were initially allowed to explore two arms (i.e., ‘familiar arms’), followed by a test phase in which the third, previously blocked arm (i.e., ‘novel arm’) was opened for exploration (Fig. 5A). As expected, WT littermate controls (n = 16) spent more time in the novel arm 95.87 ± 3.77 s compared to other arms (81.46 ± 4.76 s and 81.03 ± 3.85 s). In contrast, *Cntnap2^-/-^* mice (n = 23) did not show a preference for spending time in the novel arm (82.75 ± 4.55 s) compared to other arms (94.71 ± 4.65 s and 85.43 ± 4.74 s) (Fig. 5B). Using percentage of total exploration time spent in each arm, we found that WT littermates spent a significantly greater proportion of time in the novel arm (37.29 ± 1.58%; one-sample t-test comparing to 33.3% for equal exploration of each arm, p = 0.024), whereas *Cntnap2^-/-^* mice did not (31.45 ± 1.56%; one-sample t-test, p = 0.24). Direct comparison between genotypes showed that novel arm preference was significantly reduced in *Cntnap2^-/-^* mice relative to WT littermate controls (Welch’s t-test, p = 0.013). However, there was no sex preference for novel arm exploration (WT male: 35.94 ± 2.64%, WT female: 38.63 ± 1.79%, *Cntnap2^-/-^* male: 32.55 ± 4.12%, *Cntnap2^-/-^* female: 30.97 ± 1.45%) (Fig. 5C). These results suggest that loss of Caspr2 impairs spatial working memory, potentially through disruption of both peripheral vestibular signaling and central neural pathways.

**Figure 5.**
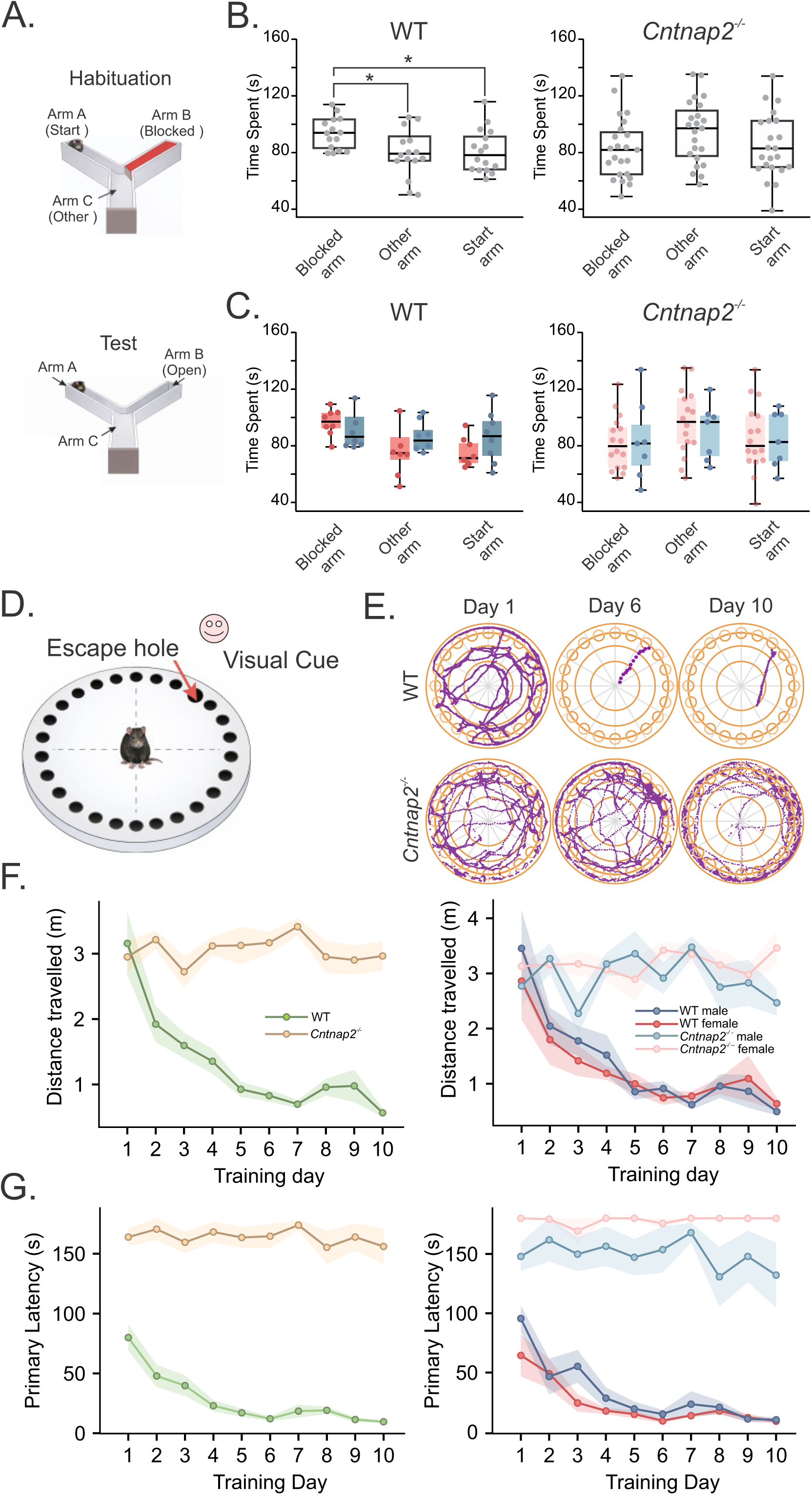
Impaired spatial working memory and spatial learning in *Cntnap2*^-/-^ mice. **(A)** Schematic of the two-trial Y-maze task. During habituation, one arm was blocked and mice explored the start arm and one familiar arm. After a 1 h retention interval, mice were returned to the maze for the test phase, during which all three arms were accessible and the previously blocked arm served as the novel arm. **(B)** WT mice preferentially explored the previously blocked, novel arm, whereas *Cntnap2^-/-^* mice distributed exploration more evenly across the three arms, consistent with impaired spatial working memory. **(C)** Novel arm preference differed by genotype but not by sex (two-way ANOVA; WT, n = 16; *Cntnap2^-/-^*, n = 23; main effect of genotype, p = 0.018; main effect of sex, p = 0.89; genotype x sex interaction, p = 0.38), so that WT mice spent significantly more time in the novel arm than expected by chance. **(D)** Schematic of the Barnes maze used to assess spatial learning. Mice were placed on a circular platform containing 20 peripheral holes, one of which led to an escape box. Visual cues surrounding the maze were used as spatial landmarks. **(E)** Representative Barnes maze trajectories from WT and *Cntnap2^-/-^* mice on training Days 1, 6, and 10. WT mice developed progressively more direct search paths across training, whereas *Cntnap2^-/-^* mice continued to show prolonged, disorganized search trajectories. **(F)** Distance traveled during Barnes maze training, shown by genotype and by sex (WT, 8 females and 8 males; *Cntnap2*^-/-^, 7 females and 7 males). WT mice showed a marked reduction in path length across training days, consistent with improved search efficiency, whereas *Cntnap2^-/-^* mice showed little change in distance traveled. **(G)** Primary latency to reach the escape hole across 10 days of Barnes maze training, shown by genotype and by sex. WT mice showed progressive improvement in latency to reach the escape hole across the 10 days of training, whereas *Cntnap2^-/-^* mice remained impaired.

### Impaired spatial learning in *Cntnap2*^-/-^ mice

We next used the Barnes maze (Fig. 5D) to assess spatial learning in *Cntnap2^-/-^* mice (7 males and 7 females) compared to WT littermate controls (8 males and 8 females). Representative tracking plots (Fig. 5E) show while WT and KO animals similarly explored the maze initially, they were clearly different in later days. WT and KO mice did not differ significantly in distance traveled on Day 1, indicating comparable initial search path length at the start of training (Welch’s t-test, p = 0.70). By the final acquisition day, *Cntnap2^-/-^* mice traveled significantly farther than WT controls before reaching the escape endpoint (2.96 ± 0.23 m vs. 0.57 ± 0.06 m, Welch’s t-test, p < 0.0001).

On average, WT mice showed a progressive reduction in path length across acquisition training (Repeated measures Welch , genotype x day interaction, F(9, 29) = 7.26, p < 0.0001), decreasing from 3.16 ± 0.48 m on Day 1 to 0.57 ± 0.06 m on Day 10. In contrast, *Cntnap2^-/-^* mice showed no reduction from Day 1 to Day 10 (Day 1: 2.95 ± 0.24 m; Day 10: 2.96 ± 0.23 m) (Fig. 5F).

WT mice showed significant improvement in learning, based on the time needed for finding the escape box (Cox proportional hazards regression model with right-censoring, hazard ratio [HR] per day = 1.21, 95% CI: 1.16-1.26, p < 0.0001). On the other hand, Cntnap2^-/-^ mice showed markedly impaired performance with few successful escapes during the 180 s time limit (43/280, 15.4% vs 315/320, 98.4% for WT controls). We found reduced learning and sex-dependent severity in the KO group (genotype x day interaction, p = 0.0007 and genotype x sex interaction, p = 0.00023). Finally, compared to sex matched WT controls, both female and male KO mice were significantly impaired, with stronger effects in females (female KO vs female WT: HR = 0.01, p < 0.0001; male KO vs male WT: HR = 0.17, p < 0.0001) (Fig. 5G). To assess memory retention, mice underwent a 1 min probe trial 3 days after the final acquisition session. Consistent with their lack of learning during the 10 day training period, Cntnap2^-/-^ mice performed poorly in the probe test, with only 14.3% (2/14 mice) visiting the former escape hole, whereas all WT mice (16/16) went directly to the previous escape location.

Together, the Y-maze and Barnes maze results indicate that loss of Caspr2 is associated with impairments in both spatial working memory and spatial learning.

## Discussion

ASD is associated with abnormalities in sensory processing, balance, motor coordination, and spatial cognition [1–3]. Although these phenotypes are often attributed to central circuit abnormalities, increasing evidence suggests that peripheral sensory dysfunction can also shape behaviors relevant to ASD [8–10]. Here, we show that disruption of *Cntnap2*, a gene associated with ASD, impairs vestibular sensory signaling and is associated with deficits in gravity sensation, balance control, and spatial learning and memory. These results identify *Cntnap2*/Caspr2 as a regulator of vestibular function and support a model in which altered vestibular input contributes to sensorimotor and spatial cognitive phenotypes in the *Cntnap2*^-/-^ model.

Recent studies identified Caspr2 at the vestibular type I hair cell – calyx synapse [38, 39] and our transcriptomic analysis shows that *Cntnap2* is expressed in vestibular sensory tissue during postnatal development. *Cntnap2* expression increases during the first postnatal month, a period during which vestibular hair cells and afferents mature [65, 66]. This developmental expression pattern suggests that Caspr2 may contribute to maturation, stabilization, or function of vestibular sensory circuits. The reduced VsEP amplitudes and prolonged P1 latencies observed in adult *Cntnap2*^-/-^ mice indicate that loss of Caspr2 produces persistent impairment of peripheral vestibular afferent signaling.

### Selective vestibular sensory dysfunction

Vestibular nerve afferents are commonly classified as regular or irregular based on the variability of their resting discharge [72]. These afferents receive input through two different synaptic mechanisms [73, 74]: a glutamatergic quantal transmission from type II hair cells [46, 75] and a specialized potassium dependent nonquantal transmission at type I hair cell – calyx synapses [43, 46, 49]. VsEPs primarily reflect synchronized activation of otolith vestibular afferents, especially irregular afferents [76] that innervate central zones of the vestibular sensory epithelium [73]. The reduced VsEP amplitudes and prolonged P1 latencies in *Cntnap2*^-/-^ mice therefore suggest impaired recruitment, synchrony, or temporal precision of vestibular afferents that innervate the otolith organs.

The mechanism by which Caspr2 loss impairs vestibular afferent signaling remains to be determined. Caspr2 organizes specialized axonal and synaptic domains and influences ion channel localization and neural excitability [33, 36, 41, 77, 78]. In the vestibular periphery, Caspr2 localization near calyx terminals raises the possibility that its loss disrupts the microenvironment of the type I hair cell – calyx synapse [38, 39]. This interpretation is supported by studies of Caspr1, a related protein whose loss alters the structure of the vestibular calyx synapse and disrupts KCNQ potassium channel localization [42]. Because nonquantal transmission depends on calyx geometry, synaptic cleft architecture, and potassium channel function [43], Caspr2 deficiency could impair structural or molecular specializations required for rapid non-quantal transmission.

### Gravity signaling, VOR, and balance

The vestibular reflex findings suggest that *Cntnap2*^-/-^ mice do not have complete vestibular failure, but instead show selective disruption of vestibular signaling. Contact righting and OCR both require detection of head orientation relative to gravity. *Cntnap2*^-/-^ mice showed delayed contact righting and reduced OCR gain, indicating impaired gravity dependent otolith signaling. In contrast, rotational VOR gain and phase were preserved across the frequencies tested. These results suggest that the canal driven VOR pathway examined here remains grossly functional, whereas gravity dependent vestibular pathways are more strongly affected.

This pattern is important because different vestibular reflexes may depend on partially distinct peripheral signaling mechanisms. VOR pathways have been linked to nonquantal transmission at type I hair cell – calyx synapses [48, 51], whereas gravity related behaviors may depend more strongly on glutamatergic transmission involving type II hair cells and bouton afferents [48]. The impaired contact righting and OCR responses in *Cntnap2*^-/-^ mice are therefore compatible with altered gravity dependent otolith signaling, potentially involving quantal glutamatergic transmission. However, Caspr2 expression in bouton terminals has not been directly established and our data do not identify the precise cellular site of dysfunction. In addition, preserved rotational VOR does not exclude central compensation or rule out abnormalities in other vestibular, cerebellar, or oculomotor pathways. The current results are therefore best interpreted as evidence for selective pathway dysfunction after *Cntnap2* deletion. Future direct ultrastructural, immunohistochemical, conditional knockout, and rescue studies will be required to determine the effect of Caspr2 loss on calyx microdomains, bouton afferents, ion channel localization, and synaptic transmission, myelination.

The balance beam findings extend this reflex phenotype into a freely moving behavioral context. This task requires integration of vestibular, proprioceptive, visual, cerebellar, and motor control signals to maintain posture during locomotion. *Cntnap2*^-/-^ mice crossed the beam in a similar amount of time as WT controls, indicating that they were able to complete the task and that their performance was not explained by gross immobility. However, they showed significantly more hindlimb slips and larger tail excursions. Because mice use the tail to generate stabilizing angular momentum during challenging locomotor tasks [70, 71], the larger tail excursions in *Cntnap2*^-/-^mice suggest greater reliance on compensatory postural strategies. This phenotype is compatible with altered vestibular contributions to balance control, including possible involvement of vestibulospinal pathways. However, balance beam performance is not exclusively vestibular, and cerebellar dysfunction, proprioceptive abnormalities, motor planning deficits, and altered central sensorimotor integration may also contribute. Thus, the balance phenotype likely reflects combined effects of peripheral vestibular dysfunction and central sensorimotor abnormalities.

### Vestibular input and spatial cognition

In addition to vestibular and balance abnormalities, *Cntnap2*^-/-^ mice showed impaired spatial working memory in the Y maze and impaired spatial learning in the Barnes maze. Vestibular input provides critical information about head motion, self-motion, and orientation relative to gravity, and these signals are integrated within hippocampal and parahippocampal circuits that support spatial navigation [15, 22–26, 28, 30]. Disruption of vestibular input can impair hippocampal spatial representations and degrade spatial learning in rodents [15, 24, 28–30]. Altered vestibular signaling in *Cntnap2*^-/-^ mice may therefore contribute to spatial cognitive deficits by reducing the reliability of sensory input needed to support hippocampal spatial coding.

The spatial behavioral findings are not easily explained by generalized immobility. In the Barnes maze, WT and *Cntnap2*^-/-^ mice did not differ significantly in distance traveled on the first acquisition day, indicating comparable initial exploration. We believe that since Caspr2 is expressed throughout the brain [33, 77, 78], the spatial deficits likely reflect interaction between altered vestibular input and intrinsic abnormalities in hippocampal, cerebellar, or cortical circuits. In addition, because *Cntnap2*^-/-^ mice showed severe impairment during Barnes maze acquisition, poor probe performance should be interpreted primarily as a consequence of impaired spatial learning rather than as isolated evidence of impaired memory retention.

Some findings also suggested sex dependent differences. Vestibular and spatial behavioral abnormalities were present across sexes, but righting latency and Barnes maze performance differed between male and female *Cntnap2*^-/-^ mice. These effects should be interpreted cautiously because the study was not designed primarily to define sex specific mechanisms and sample sizes for some behavioral assays were modest. Nevertheless, these results suggest that the behavioral consequences of *Cntnap2* deletion may be modified by sex dependent developmental or compensatory mechanisms.

In summary, our findings identify CNTNAP2/Caspr2 as a regulator of vestibular sensory signaling and reveal vestibular dysfunction in *Cntnap2^-/-^* mice. The combination of impaired vestibular nerve responses, abnormal gravity dependent reflexes, altered balance control, and spatial cognitive deficits supports a model in which disrupted vestibular input contributes to sensorimotor and cognitive phenotypes relevant to ASD. However, *Cntnap2^-/-^* mice are global knockouts, and Caspr2 is expressed in both peripheral sensory systems and central neural circuits. Therefore, the present data cannot establish that vestibular dysfunction alone causes the observed balance and spatial memory deficits. Rather, analogous to prior studies showing that peripheral somatosensory dysfunction can shape ASD related behaviors [8–10], we propose that disruption of vestibular input may alter central vestibular, cerebellar, hippocampal, and cortical pathways, thereby contributing to the ASD related phenotype in this model. These results broaden the biological framework for Cntnap2 associated neurodevelopmental phenotypes and suggest that vestibular function may be a useful domain for phenotyping and future intervention studies in ASD.

## Acknowledgements

We thank David Mohr and Corina Antonescu with the Integrated Genomics Center at Johns Hopkins University for their discussions and assistance with RNA sequencing and analysis. This work was supported by NIH-NINDS grant R01NS146406 and NIH-NIMH grant R03MH127401 to TD, by NIH-NIDCD grant R01DC019380 to SGS, by NIH-NIGMS grant R35GM156374 to LDF, by a summer Provost’s Undergraduate Research Award (PURA) from Johns Hopkins to YS, and by Lloyd B. Minor Center for Vestibular and Skull Base Sciences.

## Data availability

All data for this study are included in this published article. If needed, other information is available on request from the authors.

## Author Contributions

TD and SGS conceived the study and supervised the project. SGS wrote the initial draft of the manuscript. YS performed VsEP and balance beam tests, analyzed part of the data and wrote the related Methods and Results sections. YC performed Y-maze and Barnes maze tests, performed all data analyzed with DeepLabCut, and wrote related Methods and Results sections. DZ performed eye movement recordings, analyzed data, and wrote related Methods and Results sections. XD performed some of the Barnes maze tests. LDF analyzed RNA sequencing data and wrote related Methods and Results sections. All authors approved the submitted manuscript.

